# Not everything, not everywhere, not all at once: a study of brain-wide encoding of movement

**DOI:** 10.1101/2023.06.08.544257

**Authors:** Ziyue Aiden Wang, Susu Chen, Yi Liu, Dave Liu, Karel Svoboda, Nuo Li, Shaul Druckmann

## Abstract

Activity related to movement is found throughout sensory and motor regions of the brain. However, it remains unclear how movement-related activity is distributed across the brain and whether systematic differences exist between brain areas. Here, we analyzed movement related activity in brain-wide recordings containing more than 50,000 neurons in mice performing a decision-making task. Using multiple techniques, from markers to deep neural networks, we find that movement-related signals were pervasive across the brain, but systematically differed across areas. Movement-related activity was stronger in areas closer to the motor or sensory periphery. Delineating activity in terms of sensory- and motor-related components revealed finer scale structures of their encodings within brain areas. We further identified activity modulation that correlates with decision-making and uninstructed movement. Our work charts out a largescale map of movement encoding and provides a roadmap for dissecting different forms of movement and decision-making related encoding across multi-regional neural circuits.

## Introduction

A standard template for the function of the nervous system is the translation of sensory inputs into action^1–3^. Classical descriptions of the brain parcellate the brain into sensory and motor areas^4^. On the other hand, decades of neurophysiological recordings have found activity related to movement throughout sensory and motor regions of brain. For example, neurons in visual cortical areas are modulated by eye movement in primates ^5, 6^ and mice^7^, neurons in barrel cortex are modulated by movement of the whiskers in rodents ^8–10^, and activity related to locomotion causes widespread modulation of neural activity across multiple cortical regions^11–16^. A series of recent studies further demonstrated that movement-related signals make up a large proportion of ongoing neural activity across both sensory and motor areas^17–20^. These studies typically examined a few brain areas at a time and different studies relied on diverse behaviors and methods. There has not been a comprehensive characterization of movement-related activity across the brain in a single behavior. It thus remains unclear how movement-related activity are distributed across the brain and whether there are systematic differences between brain areas.

Moreover, the presence of motor signals by itself does not clarify the nature of such signals. For example, movement-related activity could reflect motor commands, efference copies, reafferent signals from sensory organs, mixtures of these signals, or activity related to behavioral state^21^. Recent video analysis methods that relate ongoing movements to neural activity has begun to allow for quantitative modeling of these signals^22–24^. Yet, most existing methods do not distinguish between different types of neural correlates of movement.

The ubiquity of motor signals also raised the question of how these signals influence computations performed by individual brain areas. In sensory cortical regions, movement-related activity has been shown to modulate sensory coding^12, 13, 15, 25^. But in most other brain regions, including frontal cortices, the impact of movement-related activity on neural coding is not understood. This problem is particularly pressing for interpretation of cognitive signals such as those underlying decision-making. Indeed, most cognitive tasks require animals to perform instructed overt movements to report decisions, e.g., by pressing a lever. However, animals additionally perform other movements that are not instructed. While uninstructed, in many cases these movements are correlated with the cognitive process under study^26^. For example animals may perform small movements biased towards the direction of future choice as evidence is accumulated. Indeed, signals related to accumulated evidence has been reported in muscle tensions^27^ or even in ongoing movement executions ^28, 29^. Under some conditions, animals could be trained to be perfectly still and avoid these uninstructed movements^30^, but this is not always possible, and fine movements, postural changes, and adjustments of muscle tension could be hard to detect and to suppress. This causes a potential confound between pure cognitive representations and neural activity related to uninstructed (or instructed) movements. While previous studies have highlighted motor-related signals in decision-making and motor planning areas of the brain^17, 19, 22^, most previous did not parcellate decision- and movement-related activity and compare different type encodings across brain regions. Consequently, it remains unclear how pervasive uninstructed movement signals are across the brain, and how they are related to neural activity modulated by an animal’s decision.

To answer these questions, we analyzed a comprehensive set of population recordings containing more than 50,000 neurons across the brain, including over a dozen cortical and sub-cortical structures, simultaneously with high speed video of orofacial movements of the mice performing a decision-making task^31^. We tested multiple methods to solve the challenging computational problem of relating two high-dimensional, complex time-series datasets: pixels of behavioral videos describing the movement of the animal and time-varying spike rates of recorded in specific brain regions andanalyzed the relation between movement and neural activity. We found that though movement-related signals were widespread, the predictive power of movement related signals was systematically different across areas, with increasing predictive power in areas closer to the motor or sensory periphery. We further dissected motor-related versus sensory-related signals in close relationship to the underlying anatomy and found maps of sensory and motor processing. Further, we revisited the long suggested but relatively unexplored question of uninstructed movements and their relation to decision-related signals, and were able to distinguish between neurons whose modulation is primarily movement-dependent versus others whose modulation is more purely task-contingency-dependent. The prevalence of these two types of neurons systematically differed across brain areas. Our study offers a roadmap for dissecting the relationship of movements and cognition across multi-regional neural circuits.

## Results

### Neural activity explained by movement is pervasive but differs substantially across brain areas

Populations of single units were recorded while mice performed a decision making task. (Fig. 1). In brief, mice were trained to perform directional licking (lick-left or lick-right) depending on the frequency of a series of pure tones presented to the animal (12 kHz tones instructs lick-left; 3 kHz tones instructs lick-right) to obtain water reward^32^ (Fig.1a). In between the stimulus delivery and the behavioral response, mice were required to withhold licking for 1.2 seconds. We refer to the time period in which the sensory stimulus is presented as the "sample" epoch, the period in which mice were required to respond as the "response" epoch, and the period in between as the "delay" epoch. High-speed (300 Hz) videos of the face and paws were acquired from side and bottom views, together with neural population recordings (Fig.1b). Two to five Neuropixels^33^ probes were used simultaneously to record extracellular activity in multiple regions of the mouse brain, including anterolateral motor cortex (ALM), an area critical for directional licking decisions, as well as medulla, midbrain, striatum, and thalamus (Fig.1b-d). Recording locations were registered to the Allen Common Cooordinate Framework (CCF) and thus mapped^38^ to the Allen Reference Atlas (ARA)^39^.

**Figure 1.**
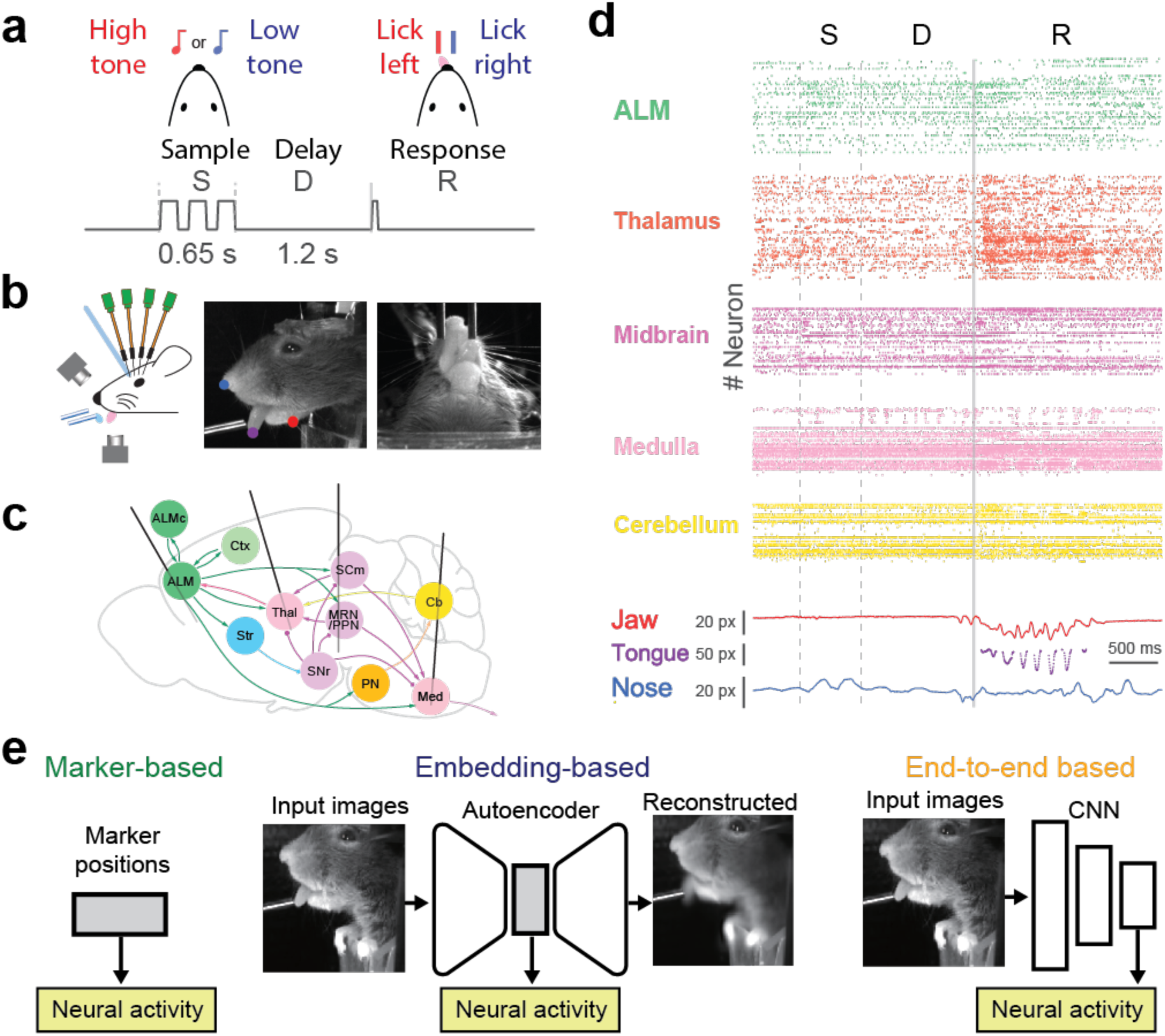
Comprehensive neural recordings in a single task and prediction of neural activity from video. **a.** Schematic of delayed response task and its epochs. **b,** schematic of simultaneous video and neural recording **c.** Schematic of brain areas and probe penetrations. **d.** Raster plot of recorded neurons (top) and traces of body part marker locations for a single trial. **e.** Schematic of the three approaches to predict neural activity from video. Left: marker-based analysis pipeline. For each video frame, each of the marker (jaw, nose or tongue) is a 2-dimensional vector representing the vertical and horizontal position. Middle: embedding-based analysis pipeline. For each frame, the embedding vector is a 16-dimensional vector. Right: end-to-end learning with deep neural network pipeline.

To examine how pervasive was neural activity related to ongoing movements, we analyzed the relation between facial and paw movements during performance of the delayed response task and related neural activity to movements using three approaches: marker-based, embedding based, and end-to-end learning. In more detail, our first approach was marker-based (Fig. 2a). We selected three markers: a nose marker, a tongue marker, and a jaw marker. DeepLabCut^40^ was used to track the two-dimensional location of the feature in each video frame. We then regressed neural activity based on the time series of the positions (Fig. 1e, left). The second approach was based on first learning a low-dimensional embedding of the videos via an autoencoder^41^, which we refer to as the embedding-based method. We trained autoencoders to reconstruct each frame through a low-dimensional bottleneck (Fig. 1e, middle). The encoder was a convolutional neural network, and the decoder was linear (see Methods). In this architecture each frame was transformed into a 16-dimensional vector, i.e., a point in the emedding space. When relating the markers and embedding space to neural activity we took a five frame window to simplify representation of movements, resulting in an 15-dimensional space for markers and an 80-dimensional space for the embedding which was then related to neural activity. The third approach used direct end-to-end learning. Instead of transforming the videos into a different representation independently of neural activity and then relating that representation to neural activity, we trained neural networks to directly predict activity from video (Fig. 1e, right, and see Methods). The first approach was the least expressive, as we manually selected three features. The second approach was more expressive in that the non-linear encoder network could learn a richer, if still low dimensional, representation. The third was the most expressive as it could make full use of the high dimensional dynamics in the video to explain activity.

**Figure 2.**
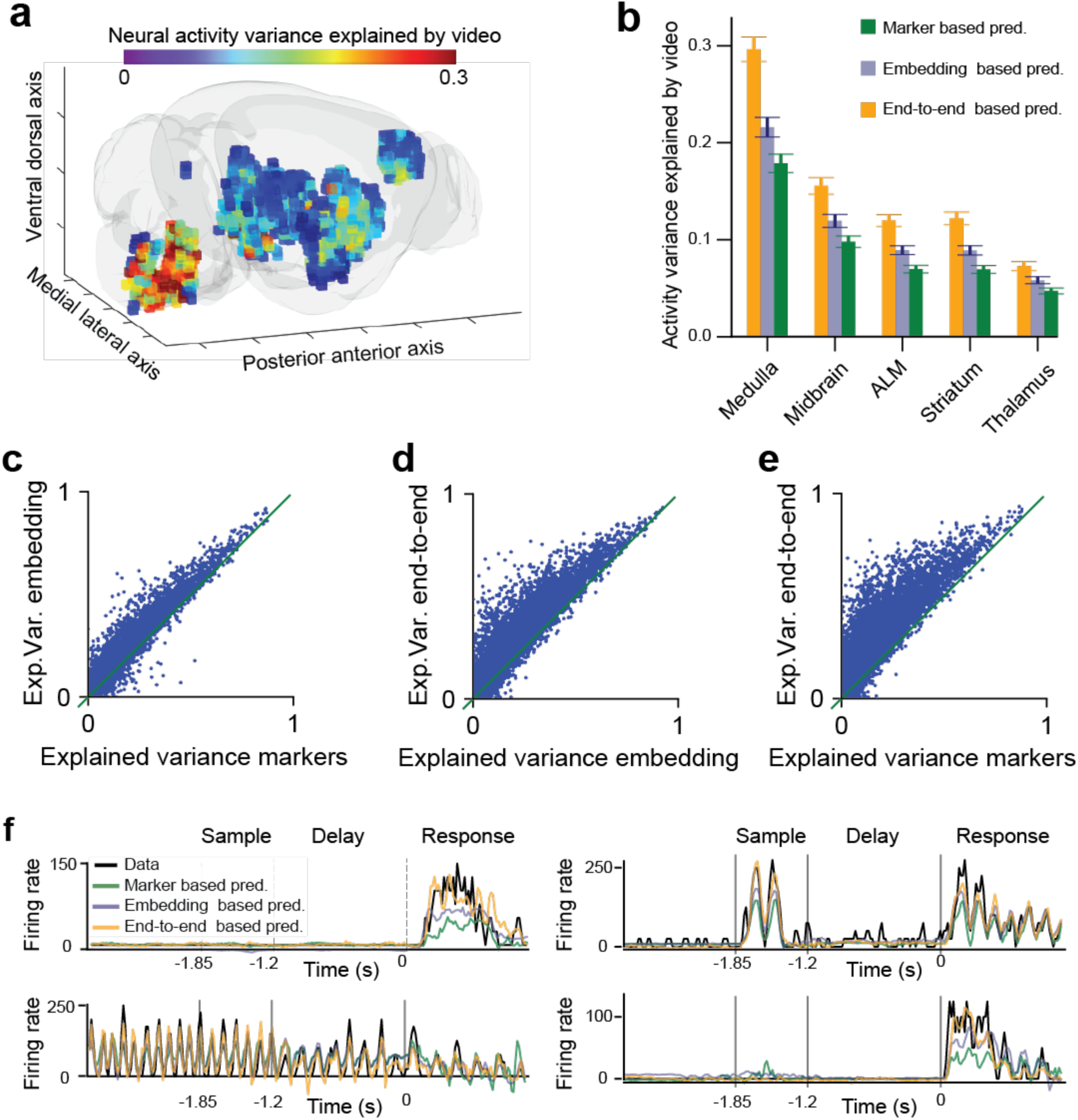
Movement encoding systematically differs across brain areas. **a.** A three dimensional spatial map of brainwide prediction of neural activity from video. Each voxel is 300 x 300 µm cubed. Color corresponds to mean variance explained by the embedding-based pipeline over all the neurons contained within each voxel. **b.** Performance of video-based prediction with neurons pooled according to brain area. Error bars correspond to standard error of the mean of session-averaged values.**c.** Comparison of single neuron explained variance between marker-based pipeline (x-axis) and embedding based-pipeline (y-axis). Each dot corresponds to a neuron **d.** Comparison of single neuron explained variance between embedding-based pipeline (x-axis) and end-to-end learning (y-axis). Each dot corresponds to a neuron **e.** Comparison of single neuron explained variance between marker-based (x-axis) pipeline (x-axis) and end-to-end learning (y-axis). Each dot corresponds to a neuron. **f.** Firing rates of four example neurons during four single trials. Firing rates are plotted (black line) overlaid with their prediction from the marker-based (green), embedding-based (blue) and end-to-end (orange) pipelines. Note negative firing rate could have been removed post-hoc but here we show raw prediction output. The explained variance for the four trajectories are the following. From top left to bottom right: medulla neuron, variance explained marker 0.09, embed 0.60, end-to-end 0.76; second: medulla neuron, variance explained marker 0.41, embed 0.66, end-to-end 0.73; third: medulla neuron, variance explained marker 0.64, embed 0.75, end-to-end 0.80; fourth: striatum neuron, variance explained marker 0.35, embed 0.48, end-to-end 0.69.

Combining neurons across animals into a registered space, our analysis recapitulated the finding that movement related activity is found across the brain (Fig. 2 a,b). Yet at the same time, it revealed clear differences across the brain in the ability of the video recordings to predict neural activity (Fig. 2 a,b) with recognizable patterns such as the dense concentration of high explained variance in the medulla (Fig. 2a). We found significant differences across brain areas (Fig. 2b). Variance explained was highest in the medulla amongst all five regions and it was significantly larger than midbrain which was at second place (medulla explained variance from the embedding method 0.216 +/- 0.06 sem, n=36 insertions compared to 0.120 +/-0.006, n=81 of midbrain, p<0.001, Mann-Whitney test, note insertions rather than sessions were used since in some sessions recordings were performed for a given brain area simultaneously across two hemispheres) an area where multiple premotor nuclei reside. Explained variance followed a logical progression with greater explained variance in areas closer to the periphery, motor, or sensory (Fig.2b).

While the ordering of brain areas in terms of predictive power was preserved across approaches, the approaches significantly differed in their predictive power (Fig. 2 c-f). We found that the more expressive models (i.e., models with larger number of parameters that can express more complex functions) better captured neural activity (Fig. 2c-e). The embedding-based approach outperformed the marker-based approach in nearly all recordings (Fig. 2c, improvement in explained variance = 23.3% +/- 0.3% sem, n=23044 neurons, p<0.001). End-to-end learning in turn outperformed the embedding based approach in nearly all recordings (Fig. 2d, improvement = 35.1% +/- 0.5%, n=23044 neurons, p<0.001) and outperformed the marker-based approach in nearly all recordings (Fig. 2e, imporovement = 66.5% +/- 1.4%, n=23044 neurons, p<0.001). Better predictive accuracy was clear when visualizing single trial predictions (Fig. 2f). However, not all neurons were predictable. These were not just low spike rate or extremely variable neurons, some neurons had highly reliable responses across trials but were poorly explained by video prediction (see Methods for definition, Fig. S1). The activity profile of some of these neurons strongly suggests that they represented the auditory cues delivered in the task (Fig. S1). Comparison across epochs suggested that a large portion of activity was correlated with licking and associated facial movements (Fig. S2)

### Dissecting putative motor and sensory neural signals

Movement-related signals are not of a single kind and could reflect motor commands, reafferent signals or combinations of the two. The high temporal information allowed by electrophsyiology and the rich dataset across multiple areas allowed us to analyze the temporal relation between neural activity and behavior (Fig. 3). We shifted the window of video frames used to predict neural activity across a range of lead or lag times. We tested time windows both from the past and in the future relative to the analyzed neural activity (Fig. 3a). For a brain area involved in producing movement, the current activity determines future movement, which will be reflected in future video frames. Thus, shifting the window of behavioral variables forward in time will yield better prediction (Fig. 3a). Conversely, if an area is sensory (e.g., proprioceptive), then current activity is related to the state of behavioral variables in the past. Thus, shifting the window of behavioral variables backward in time will yield better prediction (Fig. 3a).

**Figure 3.**
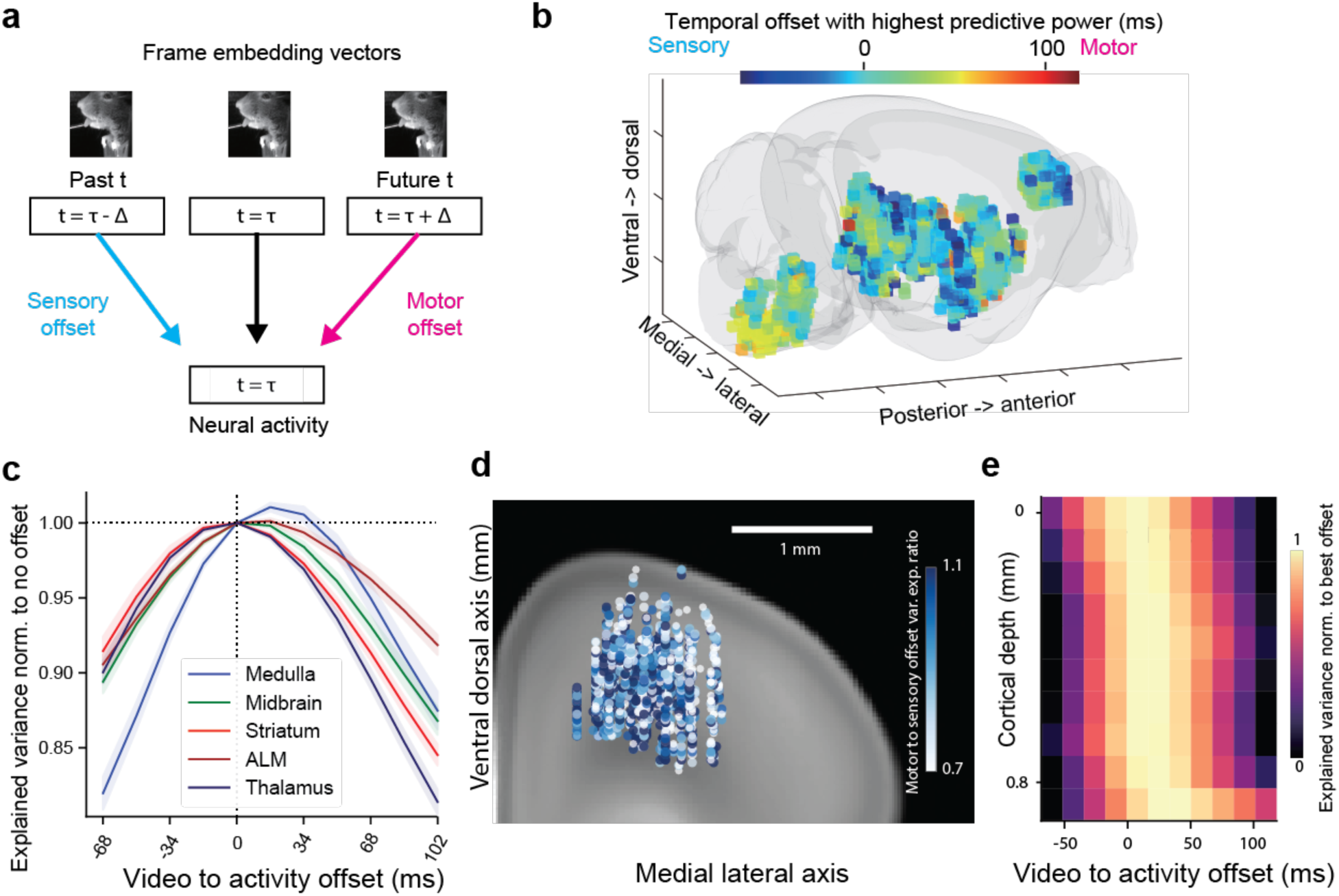
Temporal relation between neural activity and behavior. **a,** Schematic of video to activity prediction with different temporal offsets. comparison across shifted time windows. **b.** Brainwide spatial map of the best temporal offsets for each neuron. Voxels are of size 300 µm cubed. Color represents the mean best temporal offsets of all neurons within each voxel. **c.** Comparison of prediction accuracy across different temporal shifts averaged by brain area. For each area, the explained variance across different temporal shifts is shown as a line, normalized to the explained variance at zero temporal shift. Lines correspond to mean across sessions, colored by brain region; shaded area corresponds to the standard error of the mean across sessions. **d.** Heatmap of ratio of explained variance at motor offset to explained variance at sensory offset (85 ms and -51 ms). Each circle corresponds to a single ALM neuron. **e.** Comparison of explained variance across different temporal offsets as a function of cortical dept. Each row is normalized by its minimal and maximal values.

We performed this analysis in the response epoch which had the strongest relation between video and activity. We first focused on the embedding-based approach (the other approaches yielded similar results). We found clear differences across the brain (Fig. 3b) with a clear pattern of neurons in the medulla area having a strong preference for video-shifts into future time points, consistent with the known role of medulla in directly controlling movement^42^. Averaging neurons within a brain area and comparing across different brain areas, the medulla stood out for having a strong preference for video-shifts into future time points, i.e., more motor-related representations (Fig. 3c). When comparing neurons within brain areas, we found heterogeneous video-activity shift preference (Fig. 3d,e). In ALM we find a change from a more sensory-associated to a motor-associated preference (explained variance from sensory to motor offset at superficial layers changed by 6.8% +/- 0.6% sem, n=994 neurons compared to 3.9% +/- 0.6% at deep layers, n=720, p<0.001, see Methods). These findings are consistent with established sensory-motor difference of superficial layers of ALM receiving more sensory signals while deeper layers of ALM sends motor signals to midbrain and medulla^36, 43, 44^.

To test the robustness of our results we repeated these analyses with the end-to-end learning pipeline. Despite the fact that the end-to-end learning generated different and more accurate per-neuron predictions (Fig. 2) we found qualitatively similar results in terms of across area differences in sensory to motor offset explained variance (Fig. S3). Finally, we observed that the best time offset was correlated with the explained variance of the neurons (Fig. S4). Dividing the neurons into higher explained variance neurons and lower explained variance neurons we find that the best time offset of higher explained variance neurons was significantly more motor-related than those of lower explained variance neurons (time offsets: ALM 2.9 +/- 1.3 sem ms for lower explained variance neurons, n=1488 neurons, in higher explained variance neurons it was 24.2 +/- 2.7 n=270, p<0.001. Medulla 17.6 +/- 1.2, n=1322 vs 23.7 +/- 1.8, n=377, p=0.005. Midbrain 9.1 +/- 0.8, n=28254 vs 13.7 +/- 1.7, n=545,p=0.015. Thalamus -5.1 +/- 0.7, n=3941 vs 3.6 +/- 1.6, n=544, p<0.001. Striatum 0.8 +/- 0.9, n=2402 vs 7.3 +/- 2.0, n=424, p=0.003. See Methods). In other words, more motor-related neurons were better predicted by behavior videos.

Registering neurons to the CCF allowed us to look not only at the level of brain areas and cortical layers, but also to examine organization of small subcortical nuclei. Our recording dataset sample large portions of the thalamus, covering multiple thalamic nuclei. Analyzing the thalamic responses, we find sunbstantial structure (Fig. 4). Overall, explained variance was non-uniform across the thalamus (Fig. 4a, test against spatial uniformity, p<0.0001, see methods). To facilitate analysis we grouped neurons into sub-regions based on the Allen onthology. For seven of the sub-regions we had sufficient number of recorded neurons for analysis (threshold set at 100, see supplementary table S1 for definition of sub-brain regions). Explained variance based on the embedding method varied significantly among sub-regions (Fig. 4c-e). Spike rate differences did not explain differences in variance explained (spike rate differene not significant between PO, VAL, VM, VPN p>0.1, yet variance explained difference differed significantly between VAL and PO/VM/VPN). Beyond analyzing specific sub-regions we analyzed how variance explained changes over space by measuring the distribution of difference in variance explained between nearest-neighbor neurons. If variance explained changes as some smooth function over CCF space then the difference in variance explained between a neuron and its nearest-neighbors would be smaller than the difference between that neuron and a randomly selected one. We found that variance explained was significantly smaller between nearest-neighbor pairs, indicating spatially structured smooth changes in variance explained (Fig. 4f, nearest neighbor difference in explained variance smaller than shuffle control, p<1e-6). Moreover, sub-region structure was not limited to the thalamus. We found clear structure for example in the superior colliculus, SC (Fig. S5) variance explained was higher in the intermediate and deep (motor) layers which project to the medulla, but not in the superficial (sensory) layers^45^ (mean R^2^: deep mean = 0.15 ± 0.003 S.E.M., N = 1433 neurons; superficial layers = 0.04 ± 0.007 S.E.M., N = 73 neurons, KS test, p < 1e-5) and in the SC.

**Figure 4.**
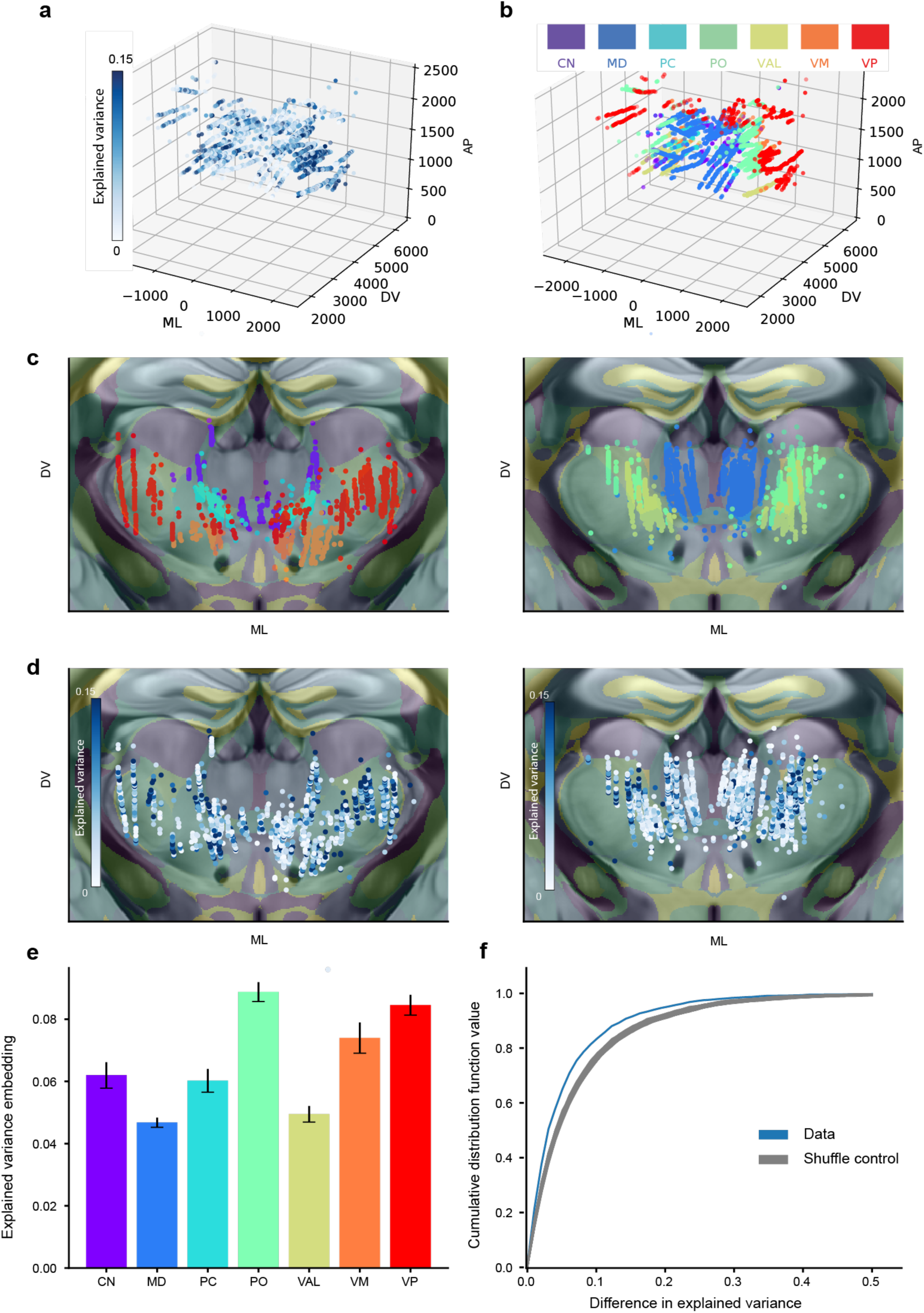
Sub-region video analysis reveals differences across thalamic nuclei. **a.** heatmap of fraction of neural activity variance explained by video prediction. Each dot corresponds to a neuron. Color reflects variance explained by embedding-based prediction **b.** Annotations for thalamic nuclei. Each dot corresponds to a neuron. Neurons were mapped to their CCF coordinates and colored according to the annotation for that coordinate. **c.** Two-dimensional projection of nuclei annotation. Each dot corresponds to a neuron. Color corresponds to nuclei annotation. Left plot shows more anterior portion of thalamus, right plot shows more posterior. Overlaid with high transparency is a map with color corresponding to Allen reference atlas annotation. **d.** Two-dimensional projection of variance explained. Each dot corresponds to a neuron. Color reflects variance explained by embedding-based prediction. Left plot shows more anterior portion of thalamus, right plot shows more posterior. Overlaid with high transparency is a map with color corresponding to Allen reference atlas annotation. **e.** Variance explained averaged by nuclei. Error bars correspond to standard error of the main. **f.** Variance explained varies non-discontinuously across space. Plot shows cumulative distribution function of difference between variance explained of each neuron and its nearest neighbor neuron. Data is in blue and one hundred repetitions of neuron-by-neuron shuffle are in gray (see methods).

### Analyzing and interpreting uninstructed movements

Are uninstructed movements related to animals’ decision? If uninstructed movements bear some relation to future behaviors then single-trial choices could be predictable directly from behavioral-videos. We trained decoders to predict choice from behavioral videos (Fig. 5). We found that before the sample epoch, predictions of trial-type where at chance (Area Under the Curve (AUC) of Receiver Operating Characteristic (ROC) was 0.51 +/- 0.06 std, n=106 sessions, prediction based on embedding method, see Methods) consistent with the lack of information regarding trial-type at this point. Second, in the sample and delay epoch, the mean AUC increased significantly (0.66 +/- 0.12 std, n=106 sessions, mean AUC ROC of the second half of sample epoch and the delay epoch). Thus, even before mice performed their explicit choice action (directional licking), uninstructed movements contain trial-type information. Third, soon after the go cue, prediction saturated at close to perfect performance (0.99 +/- 0.01 std, n=106 session, mean AUC ROC of the second half of response epoch), consistent with the choice being easily decodable from video at the time directional licking occurs. Predictive accuracy was highly variable across sessions in the sample and delay epoch (Fig. 5a-c). Future behavior was predictable from videos during the delay epoch in some sessions, but not in others. In other words, animals were highly heterogenous in the extent they exhibited non-instructed, trial-type-related movements. Variability in predictability of actions from video was smaller within mice than across mice, consistent with the notion that individual animals adopted relatively consistent degrees of non-instructed trial-type related movements. (Fig. 5d, Calinski Harabasz clustering score^46^ for clustering of within-animal points 10.45, compared to null model value of 1.02 +/- 0.37 std, higher scores correspond to stronger clustering, p<0.001, see Methods).

**Figure 5.**
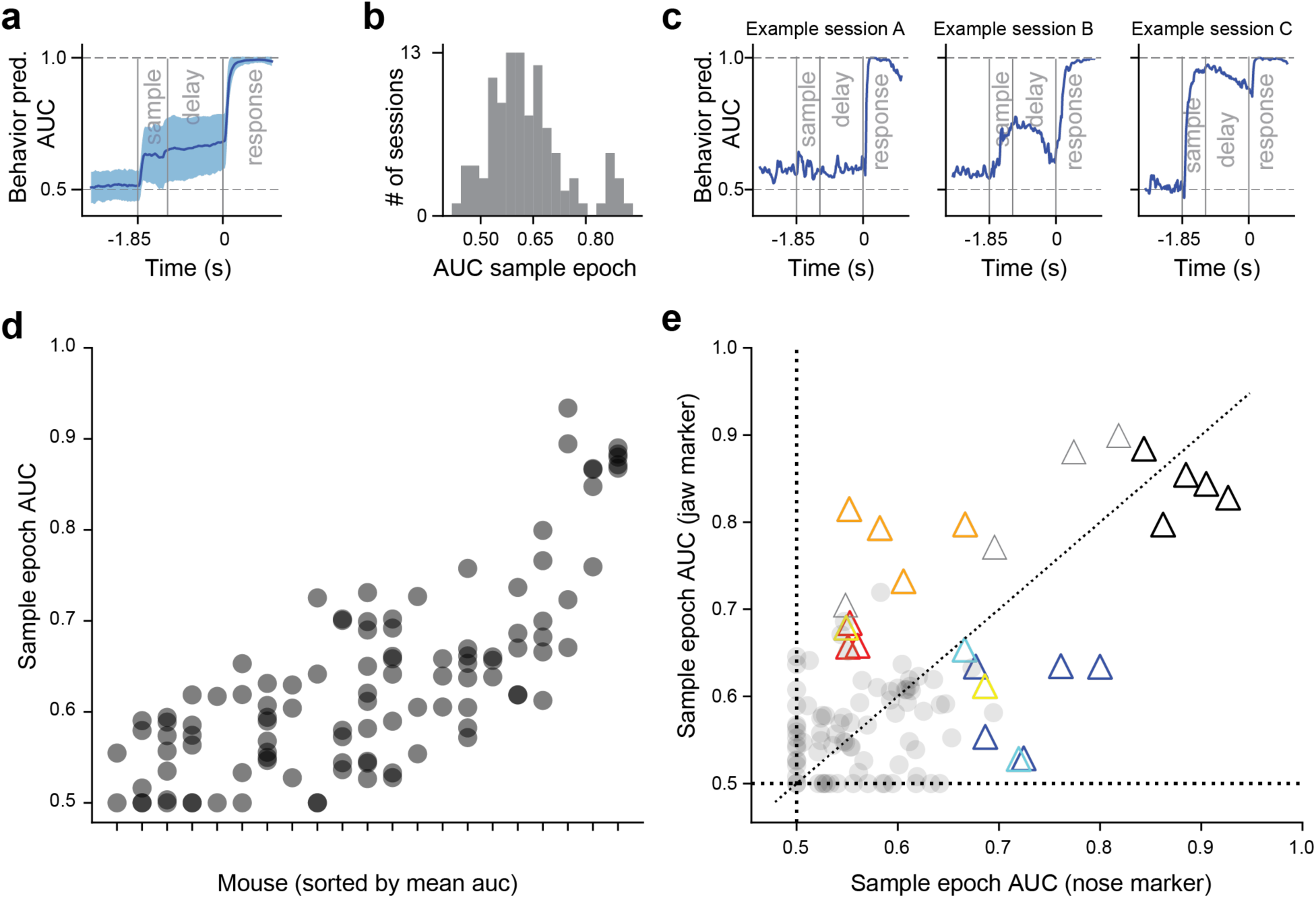
Prediction of single trial behavior directly from video. **a.** Prediction accuracy of single trial behavior from videos through embedding pipeline. Accuracy quantified through ROC AUC (y-axis). Thick line indicates across session mean. Shaded area indicates standard deviation. **b.** Histogram of prediction accuracy during the sample period across sessions. **c.** Single sessions examples. Each panel corresponds to a single session, and each session taken from a different mouse. **d.** Sample epoch prediction across all sessions and mice. Each x-axis location corresponds to an individual mouse. Each circle is a session. **e.** Prediction from single markers during the delay period. Each circle or triangle corresponds to a session. X-axis value corresponds to ROC AUC from behavioral prediction using only the nose marker. Y-axis value corresponds to ROC AUC from behavioral prediction using only the jaw marke). To allow association of sessions to mice to be visible, only sessions that passed a high predictability criterion (AUC larger than 0.6) are shown as colored traingles. The rest are shown as circles. Colors of triangles correspond to individual mouse identity.

To visualize the nature of non-instructed trial-type related movements that, we first sorted trials according to prediction confidence. We found that in highly predictable trials, mice displayed mostly stereotypical patterns of behavior, but the nature and trial-association of movements were diverse across sessions. A subset of mice tended to have more non-instructed movements in lick-left trials (Mov. S2a), while another subset had stronger movements during lick-right trials (Mov. S2b). Similarly, the movements themselves differed from mouse to mouse. For other mice, the non-instructed movements were jaw or paw movements (Mov. S2a), or only jaw movements (Mov. S2b). Furthermore, in some cases the behavior of individual mice varied across days. For example, one mouse remained static before the go cue in lick-left trials (Mov.S2b), but in the next day performed stereotypical swinging of the paw in lick-left trials (Mov.S2c).To allow more interpretable analysis of the movements that held predictive power we repeated the prediction analysis, but based on single markers, not the embedding. Overall the prediction was weaker (response period: marker: 0.88 +/- 0.01 sem, n=106, embedding: 0.96 +/- 0.00 sem, n=106, p=<1e-6, n=106) but heterogeneity of non-instructed trial-related movements was still present and variability of prediction was smaller within than between mice (Fig. 5e, Calinski Harabasz clustering score for clustering of within-animal points 12.57, compared to null model value of 1.05 +/- 0.54 std, higher scores correspond to stronger clustering, p<0.001, see Methods). For some animals, movements of the nose were more informative than movements of the jaw while in other mice jaw movement was more informative than nose movements (Fig. 5e). Finally, we found that such task-related preparatory movements during sample and delay epochs were weakly, but significantly positively correlated with the animal’s performance (r=0.33, n=106, p<0.001, Fig. S7, see methods).

### Correlating single-trial video with single-trial firing rates

Given that uninstructed movement could predict future behavior, we next explored the influence of uninstructed movement on the analysis of decision-related modulation of activity. We began by analyzing the overall dynamics of single trial predictions. Direct prediction of behavior from videos allowed us to analyze the relative timecourse of two separate predictions of behavior: a direct prediction of behavior from video and a prediction of behavior from a given area’s neural activity. Intuitively speaking, in a more motor related representation, neural activity preceds movement and therefore neural activity based prediction of behavior will rise earlier than a video based one. Comparing the time courses of prediction from neural activity and video (Fig. S6) we find that in the ventral medial medulla predictions from neural activity rose earlier and were more accurate than predictions from video during the sample period (offset in the sample epoch = (44.9 +/- 4.0) ms, p<0.001, n=28. Time offsets were calculated by the peak of the cross-correlation between the two AUC curves. AUC difference in 2nd-half sample epoch = (7.8 +/- 2.2 sem) x10^-2, p=0.002, n = 28 sessions). They then saturated at a similar level during the delay period but again rose earlier in the response period. In contrast, we found different temporal relations between video and neural prediction in the dorsal lateral medulla. Neural and video predictions had a very similar time course during the response period and video-based prediction actually outperformed neural based prediction during the delay period (AUC difference in 2nd-half sample epoch = (5.9 +/- 2.1 sem) x10^-2, p=0.012, n = 22 sessions). We note that the population level predictions were based on not-small populations. The number of neurons in the two areas was 920 from 24 recordings in ventral-medial, and 629 in 21 recordings in dorsal-lateral. In the response epoch, the offset from ventral-medial was greater than that in dorsal-lateral (mean offset in ventral medial = 44.9 +/- 4.0 sem ms, n=28, compared to 27.0 +/- 5.3 sem ms, n=22, p = 0.012), consistent with our previous finding that ventral-medial is more motor-associated than dorsal lateral.

In the previous section we trained classifiers to predict trial-type from the video embedding (Fig. 5). Next, we analyzed the relation between the single-trial output of this classifier and neural activity. When movements can predict future choice, movement encoding is conflated with choice encoding. To better dissociate choice and movement encoding we contrasted trials of the same choice based on the state of video predictors. We analyzed correct trials only and separately divided each set of trials corresponding to a choice (lick-left and lick-right trials) into two groups based on the prediction of the video-based classifier. For example, we split all correct lick-left trials into those for which the video prediction from the delay period was lick-left and those for which the video prediction was lick-right (see Methods). Intuitively speaking, trials in which the video predictor output was the same as the actual trial-type correspond to trials with more typical movements for that trial-type, whereas trials in which the video predictor output corresponded to the other trial types correspond to trials whose movement pattern was different than typical and often more similar to the movement pattern typical to the other trial-type (Fig. 6a). This analysis divides correct trials into four groups: lick-right trials with lick-right video predictor output, which we will refer to as R-vR trials, lick-right trials with lick-left video predictor output, which we will refer to as R-vL trials, lick-left trials with lick-left video predictor outputs L-vL and lick-left trials with lick-right video predictor output (L-vR). Considering non-instructed movements, we found for example that marker position (jaw height) was significantly different across same trial-type groups with different video prediction outputs in most sessions (Fig. 6b, 56 sessions out of 80 were significant at p < 0.05 for R-vR vs. R-vL, and 56 sessions for L-vR vs. L-vL. Note only sessions with moderate and higher behavioral predictability were chosen for this analysis, defined as AUC > 0.6, see Methods). This was not surprising given that these groups were defined based on video output, but it is not a given since the prediction was performed on held-out data and on the embedding representation of trials.

**Figure 6.**
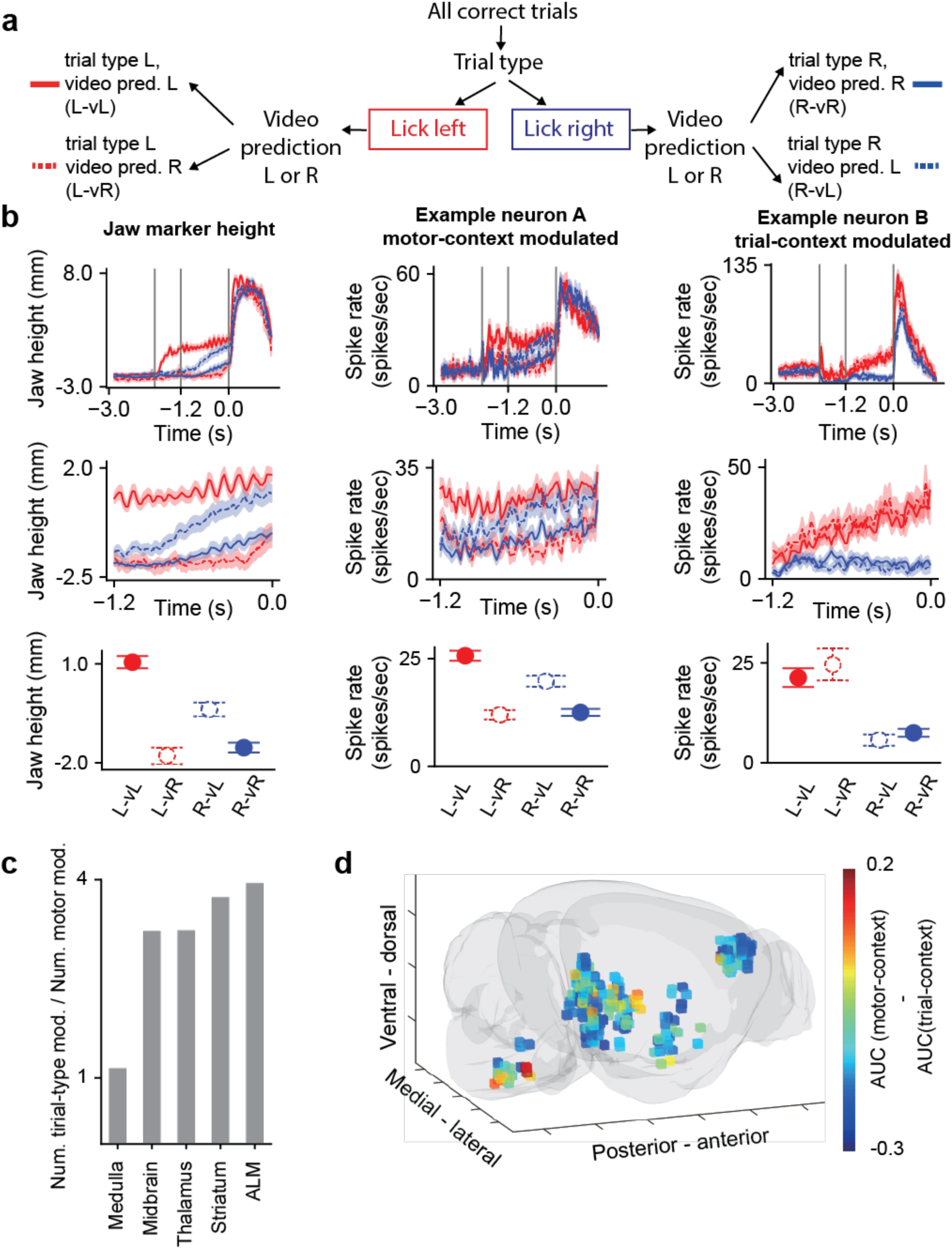
Joint analysis of single-trial video state and single-trial firing rates teases apart motor and direct trial-type related modulation. **a.** Schematic of analysis. Correct trials were split into lick left and lick right trials. Each of these sets of trials corresponding to a trial-type was further broken into two groups based on the prediction of behavior from video, yielding four groups of trials corresponding to the trial type contingency and value of single trial video prediction (see Methods). **b.** Analysis schematized in a applied to two example neurons (center, right) and jaw marker (left). <u>Left:</u> Jaw marker position (height) during entire trial (top) and zoomed in on delay epoch (middle). Lines corresponds to the mean of jaw height across the four trial groups. Color indicates trial type and line style (solid or dashed) corresponds to the video prediction based contingency. Shaded area corresponds to the standard error of the mean across trials. Center: same data as in top, zoomed in on the delay epoch. Bottom: mean jaw height during the delay epoch split into the trial type and video prediction groups. Color indicates trial type, line style type (solid or dashed) indicates video prediction contingency. Error bars correspond to the standard error of the mean across trials. <u>Middle:</u> firing rate of example neuron analyzed according to groups defined in a. Top: Firing rate of example neuron during the delay period divided into the same four groups as a. Color indicates trial type, line style indicates video prediction contingency. Center: same data as in top, zoomed in on the delay epoch. Bottom: average firing rate during the delay period split into the trial type and video prediction groups. Color indicates trial type, line style type (solid or dashed) indicates video prediction contingency. Error bars correspond to the standard error of the mean across trials. Neuron is modulated mainly by motor context <u>Right:</u> same format as in middle column for a different example neuron that is modulated mostly by trial-type context. **c.** Ratio between number of trial-type neurons, and the number of motor-type neurons, in each of the five brain regions. Neurons were classified into trial-context or motor-context modulated according to the differences across the four groups defined in a (see Methods). **d,** Brain wide spatial map of the difference between the motor-type classification AUC and the trial-type classification AUC (see Methods). Each dot is a neuron. Warm color represents motor-context modulated neurons, and cold color represents trial-type context modulated neurons.

We then analyzed each neuron for differences of spike rates across these four groups of trials. We found that some neurons’ spike rates did not change across the different video prediction output groups (R-vR vs. R-vL and L-vR vs. L-vL) but was strongly modulated by trial type (R-vR vs. L-vR and R-vL vs. L-vL). We refer to these as trial-context modulated neurons (Fig. 6b). In contrast, other neurons’ spike rate was strongly modulated by the grouping of video output even when considered within the same trial-type (R-vR vs. R-vL and L-vR vs. L-vL). We refer to those as motor-context modulated neurons (Fig. 6b). We note that some neurons were modulated by both contexts.

The relative proportion of trial-context modulated and motor-context modulated neurons varied across brain areas (Fig. 6c). In ALM we found 252 out of 283 neurons were trial-context modulated and 55 were motor-context modulated (24 neurons fell into both categories, not small neuron number was due to high single neuron signal-to-noise criteria, see methods). In contrast, among 131 medulla neurons 78 were trial-type and 71 were motor-type. The proportion of trial-type to motor-type modulation varied significantly between ALM and medulla (ALM 4.3, medula 1.1, p<0.001, binomial test) In other words, ALM neurons were more likely to be trial-type neurons than medulla neurons. This is consistent with the established role of ALM in short term memory based decision making tasks^34, 36^ and that of medulla with motor control^47^. Analyzing trial-context vs. motor-context modulation spatially, we find that across the brain the relative proportion was non uniform (Fig. 6d, defined as the average difference in AUC between the two contexts, test against uniform density, p <0.001). In summary, our analysis of the relation between video-based behavior prediction and spike rates allowed us to better disentangle neural coding of movement from decision-related activity, which revealed clear differences in encoding across different brain areas.

## Discussion

We analyzed movement related activity across the brain during a decision-making task. We found that animals typically, but not always, display uninstructed movements in a trial-type dependent manner, in addition to instructed movements. Movement-related signals were pervasive across the brain, but systematically differed across areas, with stronger movement-dependent activity in areas closer to the motor or sensory periphery. Our method further revealed structure of sensory versus motor processing within brain regions. Finally, we used single trial variance in non-instructed movements to distinguish between neurons that likely encode task-relevant information primarily due to coding of movements versus neurons whose trial-type encoding was robust but not dependent on the movements.

At a technical level, we show that high-speed video can be successfully related to population activity and that both video data and neural population are rich enough in signal-to-noise that powerful non-linear models, never before tested on this scale of data, yielded superior predictions of variance than more linear approaches.

Recent studies have emphasized ubiquitous movement related activity across the brain. While our findings are in line with these reports, our comprehensive dataset across the brain and analysis approaches were able to pull apart robust differences in the prevalence of movement related information both across brain regions and within individual brain regions. Movement-related activity could reflect motor commands, efference copies and reafferent signals from sensory ogans. Our analysis demonstrates that these types of movement signals can be dissected, and that they systematically differed across brain areas.

Importantly, combining this analysis of movement coding with detailed anatomical information for the recorded neurons was critical to reveal additional structure of sensory vs. motor processing. Within frontal cortical regions, we were able to uncover the transition from sensory to motor representations from superficial to deeper cortical layers. In the thalamus and midbrain, we were able to uncover previously reported sub-regional distinctions. Our results therefore hold promise for developing more fine-grained understanding of how distinct types of movement related information is distibuted in multi-regional circuits beyond its general ubiquity. Combining this approach with perturbation experiments could further elucidate the nature of movement-related signals and its relationship to behavior.

We found that neurons with a video-activity prediction delay consistent with motor signals, rather than a sensory reafference delay, had higher explained variance. Our interpretation is that though high speed videos contain information regarding both movement and reafference, it is likely that the relation between motor command and video is simpler than the relation between reafferance and video, and thus is more robustly captured by models in the challenging regime of signal to noise ratio of prediction of sub-single trial firing rates. More generally, we expect that different types of movements would vary in their ease of representation by models such as artificial neural networks. Future efforts are required to understand whether increasingly powerful non-linear approaches, which we have shown are possible to use given the amount of data available, might saturate out these effects or whether more explicit normalization might be required.

While most animals performed some type of uninstructed movement, they differed substantially in the degree to which these movements were dependent on trial-type. Intriguingly, we find a weak, but significant, dependence of task performance across sessions on the existence of trial-type dependent uninstructed movements, measured through classifiers’ ability to predict action at the response period from videos during the sample and delay period. Holding information in short-term memory during the delay period is known to be challenging for animals and requires extensive training. In principle, animals could assist the difficulty of maintaining information during a delay period by relying on an encoding based on different body position, either explicitly sensed during the switch to action execution or passively sensed through proprioceptive information. For multiple mice this was unlikely the primary strategy, as there were little if any uninstructed movements (and our classifiers were unable to generate useful predictions). Nonetheless, a clearer understanding of these issues would greatly benefit from the ability to modulate the degree of single trial movements in a controlled manner, either by comparing animals trained to not perform uninstrucuted movements to animals free to use uninstructed movements or by perturbation of the motor periphery to allow direct within-session comparisons of task performance across different degree of non-instructed movements. In addition, it would be interesting to test whether in more complex tasks, e.g., parameteric short-term memory tasks, context-dependent uninstructed movements are more or less informative of task condition.

When animals perform uninstructed movements that are dissimilar across trials of different types (e.g., lick-left trials vs. lick-right trials), modulation of neural activity due to task features, such as a memory of the auditory cue or planning of the future instructed movement become entangled with the encoding of the uninstructed movements. By constructing single-trial predictors of future behavior directly from video embeddings, we were able to encapsulate specific uninstructed movements that are correlated with trial contingencies. Uninstructed movements, like all movement, are not identical from trial to trial. We utilized the single-trial variability in these specific uninstructed but trial-related movements to distinguish neurons whose activity variance was explained by this variability, suggesting that these neurons encode trial-type information only indirectly through the uninstructed movements (Fig. 6b middle). These stand in contrast to neurons that encode trial-type and whose activity did not differ across uninstructed movements, which suggest they more directly reflect decision formation or maintenance of a motor plan (Fig. 6b right). Since preventing animals from performing uninstructed movements would likely be very difficult, we believe this form of video based prediction dissection of modulation will be broadly useful to isolate neural computations that give rise to cognitive signals. In particular, the question of how do uninstructed movements and their encoding develop across learning in parallel to more direct encoding of task features is an important open question.

In summary our study deepens our understanding of the encoding of movement across the brain, validates the utility of data-driven modeling of simultaneous high speed video recordings and neural population recordings.

## Methods

### Data collection and preprocessing

We analyzed data obtained from Neuropixels probes^33^ used to record extracellular activity in multiple regions of the mouse brain. To perform spike sorting, we used Kilosort^48^. We then binned spikes into firing rates with a bin width of 40 miliseconds and a stride of 20 miliseconds.

In addition to neural activity, we recorded high-speed (300Hz) multi-view video of the face, paws, and body of the mouse using CCD cameras. We used DeepLabCut ^40^ to obtain the positions of the jaw, paw, and tongue. We refer to these positions as markers. We manually labeled about 2800 frames, and trained the model using this software, and we used the same model across all sessions. Though these markers were mostly reliable, we found outliers that harmed the prediction of firing rates or animal behavior. We identified outliers by a five sigma threshold on velocity across frames and imputed outliers from nearby frames. When the tongue was occluded while it was in the mouth we set the tongue position to its mean value.

### Convolutional auto-encoder

The architecture of the convolutional auto-encoder we used similar to BehaveNet described by E. Batty et al. ^22^. The encoder was composed of two residual blocks^49^ and two fully connected layers. Each residual block is composed of four convolutional layers. The input image is resized into a 120x112 matrix. The output of the last convolutional layer is a vector with a length to be 288. This output was then processed by the two fully connected layers yielding the output of the encoder, the embedding vector, with a length to be 16. The decoder is a fully connected layer.

We trained a session specific auto-encoder. However, we also verified that one can train a session-independent encoder with session-dependent decoders. We note that decoders had to be session-dependent due to differences in overall position of the mouse, experimental components and background. In other words, during training, all frames are fed into the same encoder, but will then go to different decoders depending on which session they were extracted from. We verified that training a session-independent auto-encoder with 40 sessions, yielded similar performance and qualitatively similar analysis results to a session-dependent decoder. We also verified that the encoder can then generalize to sessions that the encoder has never seen before. This technique can be useful when one wants to train a session non-specific encoder for a very large number of sessions.

### Predicting neural activity

To predict firing rates at time t, we used embedding vectors from time steps t-34 ms, t-17 ms, t ms, t+17 ms, and t+34 ms. Thus the total dimensionality of the feature space used for embedding based prediction was 80 (the 5 neighbors times the 16 embedding dimensions). When analyzing the time offset between the embdding vectors and the firing rates, we shifted the time window to the past or to the future. For example, if the offset is +17 ms, we use the time window t-17 ms, t ms, t+17 ms, t+34 ms, and t+51 ms, to predict the firing rate at time t.

We predicted neuron spike rates using L2 regularized linear regression The value of the regularization was determined through ten fold nested cross-validation and for simplicity was maintained across time points.

### End-to-end learning framework

In the end-to-end learning framework we trained deep neural networks to directly predict neural firing rates. For each session with each brain region, we train a neural network. The network was composed of three residual blocks^49^. Each residual block was composed of four convolutional layers, and each convolutional layer is followed by a 2-dimensional batch normalization with epsilon 1 x10^-5 and momentum to be 0.1. The output of the first residual block had 16 channels, and the output of the other two blocks had 32 channels. Each residual block was followed by a 2-dimensional max pooling with a kernel size of 4 and a stride of 4. After the four residual blocks and the max poolings, the output was a vector of length 100. A linear layer was connected to the end of the last residual block. In the end-to-end learning models in Fig 2, the output was a vector whose length is equal to the number of neurons to be predicted. When considering different time offsets (Fig. S4) to speed up training we trained one network with an output of size temporal offset number times number of neurons to be predicted. In our case we used 11 different time offsets. Note that despite using one network to predict all time offsets, the predictive performance was superior to that of the embedding framework.

### Prediction of behavior

To ameliorate across session differences when predicting behavior from neural activity we first performed PCA analysis on sessions and took the top 16 PCs, to equate the size of the predictor across sessions. If a session had less than 16 neurons, we used all neurons instead of the PCs. We used ten-fold nested cross validation to estimate regularization hyperparameters.

### Defining trial-context modulated neurons and motor-context modualted neurons

We performed this analysis on correct trials only For each trial we predict behavior with the video based predictor and label the trial accordingly. We kept 80 out of 106 sessions that had high enough decodability.

To determine the modulation context of a neuron, we used the mean firing rate across an epoch to predict two single-trial variables. First we predicted the trial-type from the firing rate. Second, we used the single-trial video predictions to split all correct trial into those predicted from videos to be lick-left and lick-right trials. We then trained a decoder to predict those labels from firing rates. We label a neuron trial-context modulated if the AUC of the first decoder was higher than 0.65 and label a neuron motor-context modulated if the AUC of the second decoder was higher than 0.65. Note that a neuron can have high AUC for both predictions and thus be labeled both trial-type and motor-type.

To test the difference in prevalence of motor-context and trial-type context neuron ratios in ALM and medulla (Fig. 6h and Fig. S6h), we performed a Bernoulli t-test. We set the number of successes to be the number of trial-type neurons; set the number of “draws” to be the total number of relevant ALM neurons and set the hypothesized probability of success as the ratio between trial-type medulla neurons and the total number of relevant medulla neurons. We used p<0.001.

### Explained variance

When evaluating spike rate predictions, for each neuron *n*, we concatenate the spike rate across all trials in a single epoch into a one-dimensional vector *f*(*n*,⋅) = (*f*(*n*, 1), … *f*(*n*, *I*), where *I* is the total number of time points across all trials (or just trials if a single time-bin is considered). Similarly, we get a single vector *f*_*p*_(*n*,⋅) = (*f*_*p*_(*n*, 1), … *f*_*p*_(*n*, *I*)/ for the prediction. The explained variance of the single neuron is then

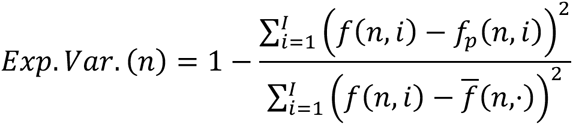

where 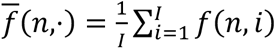

We separate trials into a train and test set using a 80/20 ratio. We note that if a model overfits on a training set the explained variance may be negative in the test set, which means the performance of the model is worse than taking the mean of the whole dataset. When using embedding vectors to predict firing rates, only 7% of the neurons had a negative explained variance. When calculating means we set the value of neurons with negative explained variance to zero as the absolute value becomes unreliable.

We analyzed the spatial smootheness of explained variance by first finding for each neuron in the analysis its nearest neighbor in space. We then calculated the difference in explained variance between the neuron and its nearest neighbor. We compared the distribution of these differences to 100 repetitions of a shuffle analysis. In each of these repetitions we randomly permuted the spatial locations across neurons, found nearest neighbor pairs (shuffling spatial location will yield in general different pairs of nearest neighbors) and calculated the distribution of differences in explained variance between nearest neighbors.

When comparing the explained variance with different time offsets (Fig. 3, Fig. 4, Fig. S4, Fig. S5), we filter out neurons with low explained variance (less than 0.02). Different choices of this threshold around similar values did not qualitatively change results.

### Calinski-Harabasz clustering score

The Calinski-Harabasz index, also known as the Variance Ratio Criterion, is the ratio of the sum of between-clusters dispersion of a measured feature to the inter-cluster dispersion for all clusters. Higher scores indicate stronger clustering. To obtain a reference null distribution for the Calinski-Harabasz score is significantly large, we randomly assign the labels (mice) to each data point, and then calculate the score, repeating the random process 1000 times. There are two inputs of Calinski-Harabasz score.

We used this clustering score two times. In Fig. 5d, we take the AUC from the embedding framework as features and mice identity as clutser labels. In Fig. 5e, AUCs from the jaw and the nose marker were used as features, and mice identity was used as cluster labels. We note that we used only sessions where the trial type was predictable from either jaw or nose (AUC equal to or higher than 0.65) and only mice that had two or more of such predictable sessions.

### Comparison performance of two prediction frameworks

We used a summary statistic referred to as bias for the comparison of two prediction frameworks. Given two sets of values *x* = (*x*_1_, … *x*_*N*_) and *y* = (*y*_1_, … *y*_*N*_), obtained from corresponding entities, e.g,. two values of explained variance based on two different frameworks on *the same neuron*, to compare the values between the two sets (frameworks) the bias, b, is defined as:

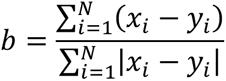

The bias b takes values between -1 and 1. When b=1, *x*_*i*_ ≥ *y*_*i*_ ∀*i*, and *y*_*i*_ ≥ *x*_*i*_ ∀*i* for b=-1. When b=0, it means the two sets of values are balanced. If the two models are equivalent the probability that an entity, i, has a higher value in one or the other is half and statistical significance can in principle be derived by integrating a binomial distribution for the number of observed entities with a higher score in framework one to obtain the probability of observing it by chance. Due to the large neuron number we approximated the binomial distribution by a normal random variable.

### Analyzing motor-sensory differences

To analyze whether two groups of neurons (group A and group B) had different statistics of motor-sensory difference we used Welch’s t-test^2^ a test designed for unequal population variance. For a given group X (X = A or B), we divided each group X into *N_X_* subgroups defined by the *N*_*X*_ sessions, and discarded sessions with less than ten neurons. For each session, we calculate the mean motor-sensory difference across all the neurons in that session. Thus, we end up with *N*_*X*_ numbers of group X, we call them *M*_*X*_ = *M*_*X*_(1), …, *M*_*X*_(*N*_*X*_). Then, we perform Welch’s t-test based on *M*_*A*_and *M*_*B*_. We perform Welch’s t-test^50^ using SciPy^51^ in python, which does not assume equal population variance.

For the comparison of video prediction explained variance and video prediction time offsets in Fig. S6 we first divided neurons in each brain region into half based on the explained variance of the neurons. We then tested the two halves for a difference in the time offset that generated the highest explained variance by Welch’s t-test.

## Acknowledgements

We thank M.N. Economo, J. Cohen, C. Poo, Chen F., Kang, B. and Kurgyis, B. for comments on this manuscript. This work was funded by the Simons Collaboration on the Global Brain (K.S., N.L. and S.D.), the NIH (NS113110 S.D., N.L. and EB028171 S.D.), McKnight foundation (N.L. and S.D.), the Sloan foundation (S.D.) and the Howard Hughes Medical Institute (K.S.).

## Supplementary Material

### Session-independent auto-encoder

We have described the session-independent auto-encoder in the method section. Here, we show the results of this method. For each of the 22 mice, we take 1 or 2 sessions to the training set, and thus we totally take 40 sessions to the training set, and we call the other 60 sessions the hold-out set. Then, we use the embedding vectors to predict the firing rates through ridge regression, in the same way, we did for the embedding from the session-dependent version. In both training set and hold-out set, the session-independent auto-encoder’s performance is much better than that of using the markers (Fig.S8a,d in the training sessions exp. var. difference 18.2% +/- 1.2% sem, n=122 recordings, p<0.001. In the hold-out sessions difference 16.5% +/- 1.1% sem, n=192 recordings, p<0.001), though it’s slightly worse than that of the session-dependent (Fig.S8b,e,c,f, in the training sessions exp. var. difference 3.6% +/- 1.1% sem, n=122 recordings, p<0.001. In the hold-out sessions difference 12.8% +/- 3.0% sem, n=192 recordings, p<0.001). This result demonstrates the encoder learns the semantic information of the mice behavior, and can generate to the sessions that the model hasn’t seen in the training process. This method also helps researchers get embedding vectors in a very efficient way, as they will only need to train one model for all the sessions.

### Thalamic sub-regions

Thalamic sub-regions were defined according to the Allen reference atlas annotations. The association between the labels used in figure 4 and annotations is given in the following table:

| Label | Associated annotations |
| --- | --- |
| CN | Central lateral nucleus; Central medial nucleus |
| MD | Mediodorsal nucleus of thalamus |
| PC | Paracentral nucleus |
| PO | Posterior complex of the thalamus |
| VAL | Ventral anterior-lateral complex of the thalamus |
| VM | Ventral medial nucleus of the thalamus |
| VP | Ventral posterolateral nucleus of the thalamus; Ventral posterolateral nucleus of the thalamus, parvicellular part; Ventral posteromedial nucleus of the thalamus; Ventral posteromedial nucleus of the thalamus, parvicellular part |

## Supplementary Figures

**Figure S1.**
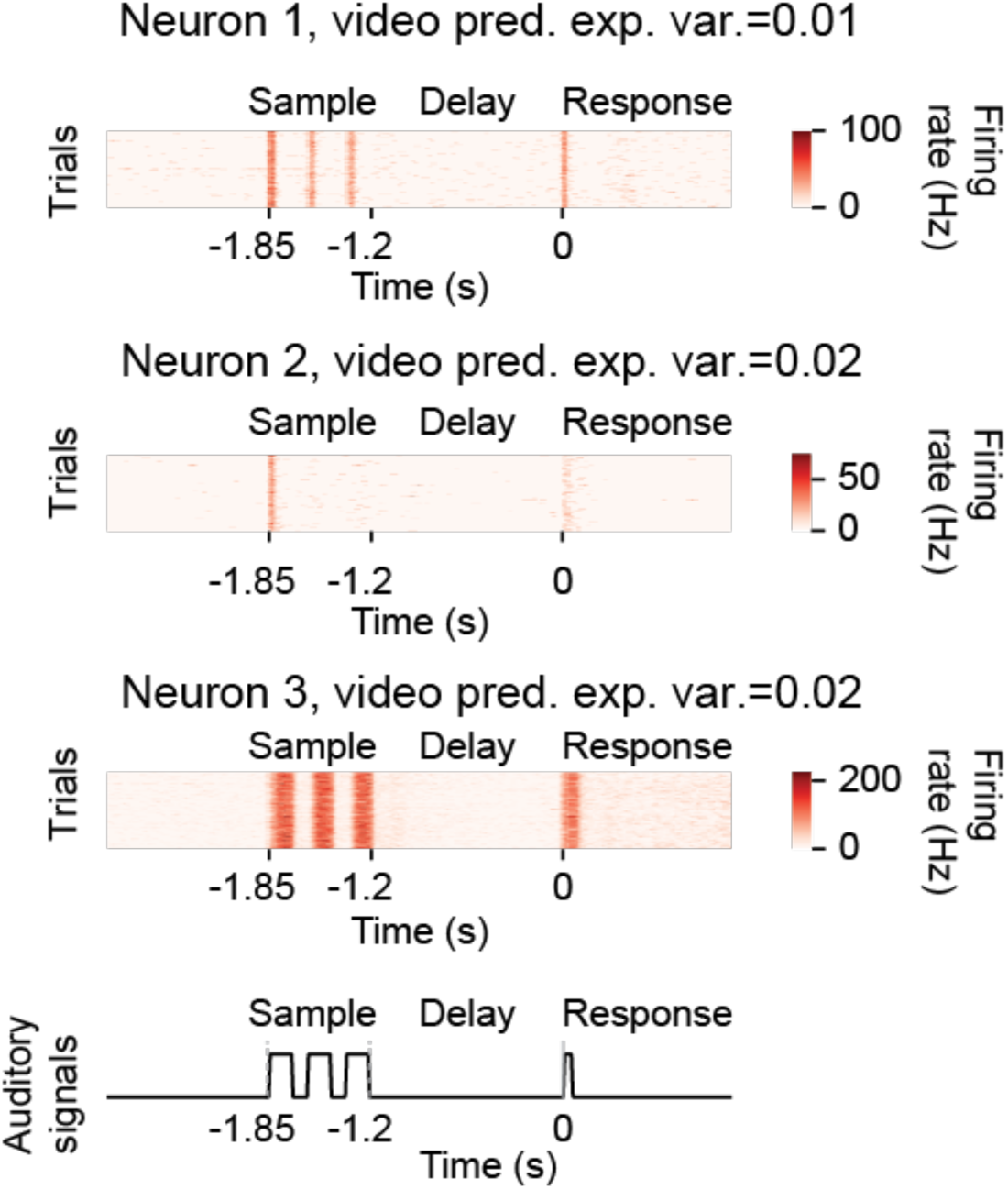
Neurons with reliable responses but whose activity is not well predicted from video. Heatmap of firing rate as a function of time. Each row corresponds to a single trial. Top neuron is a midbrain neuron, middle neuron is a striatum neuron, bottom neuron is a thalamic neuron. Bottom row shows schematic for the presented auditory cues.

**Figure S2.**
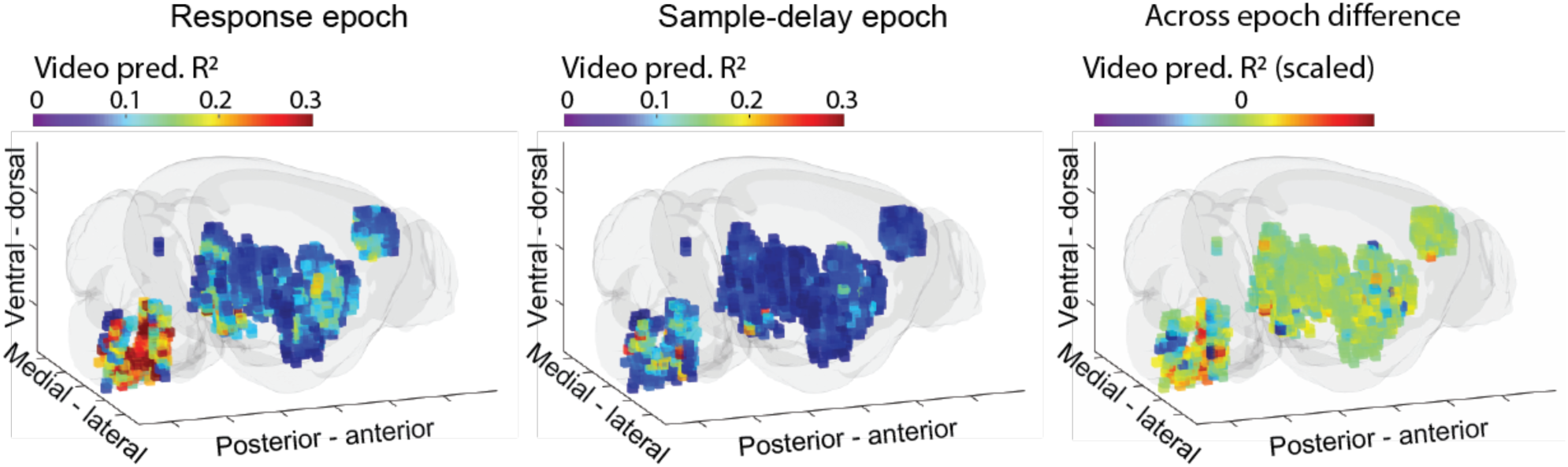
Voxel level explained variance separately per epch. Response epoch with rich licking movements (left) sample/delay epochs (middle) and difference between the response to sample/delay epochs, scaled such as the average value was zero to allow better visualization of differences from brainwide average difference.

**Figure S3.**
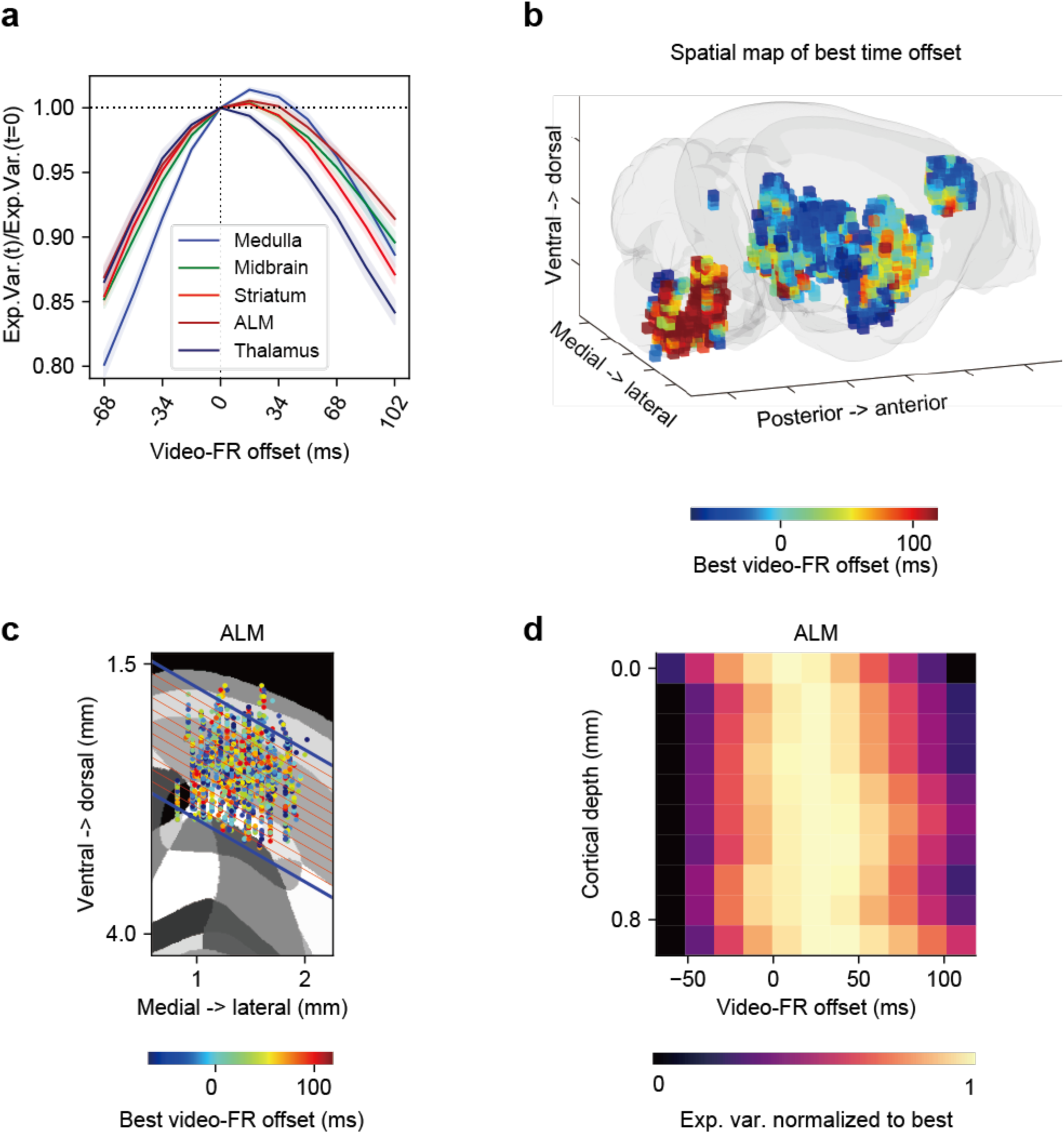
Video activity time offset analysis with end-to-end training. **a,** Comparison of prediction accuracy across different temporal shifts. For each brain region, the explained variance across different temporal shifts is shown as a line, normalized to the explained variance at zero temporal shift. Lines correspond to mean across sessions, colored by brain region; shaded area corresponds to the standard error of the mean across sessions. **b,** A 3d spatial map of the optimal time offsets across neurons from the five brain regions. Each voxel is of size 300 µm cubed. Color corresponds to the best optimal time offset for indidivudal neurons averaged across all neurons within the voxel. **c,** Single ALM neurons plotted on top of ARA from layer-2 to layer-5. Colors correspond to the time offset that gives the highest explained variance for each neuron. **d,** The heatmap of the explained variance of different time offsets, along the direction perpendicular to the direction of the red lines in c. Each row corresponds to a line in c. Each row is normalized by its minimal and maximal values.

**Figure S4.**
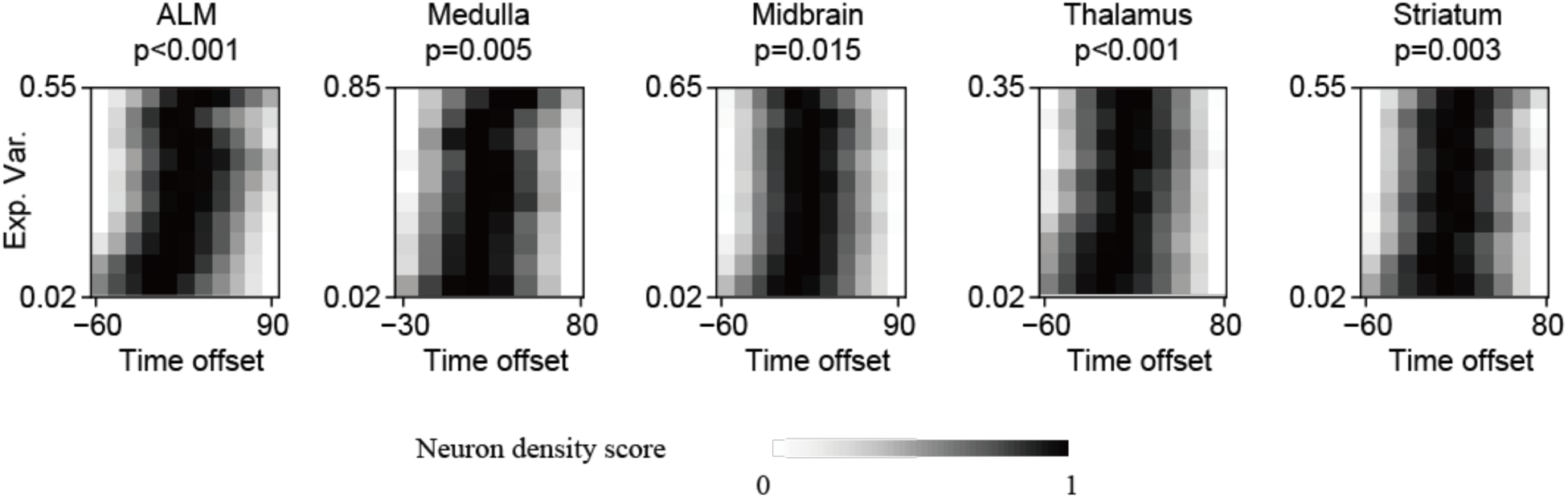
Relation between peak explained variance and video activity time offset. The peak explained variance for each neuron is the highest explained variance among the 11 time offsets tested. Neurons were first binned according to their explained variance. Brightness represent density of neurons’ optimal time offset calculated with a soft-max operation (see methods). In brief, for each neuron, its best time offset (the time offset with the highest exp. var.) will earn the highest score, 10; the second time offset will earn a score of 9, etc. Then we get the 2d histograms by pooling across neurons within that explained variance bin and calculating the density of the scores. Finally, we normalize each row of the 2d histogram.

**Figure S5.**
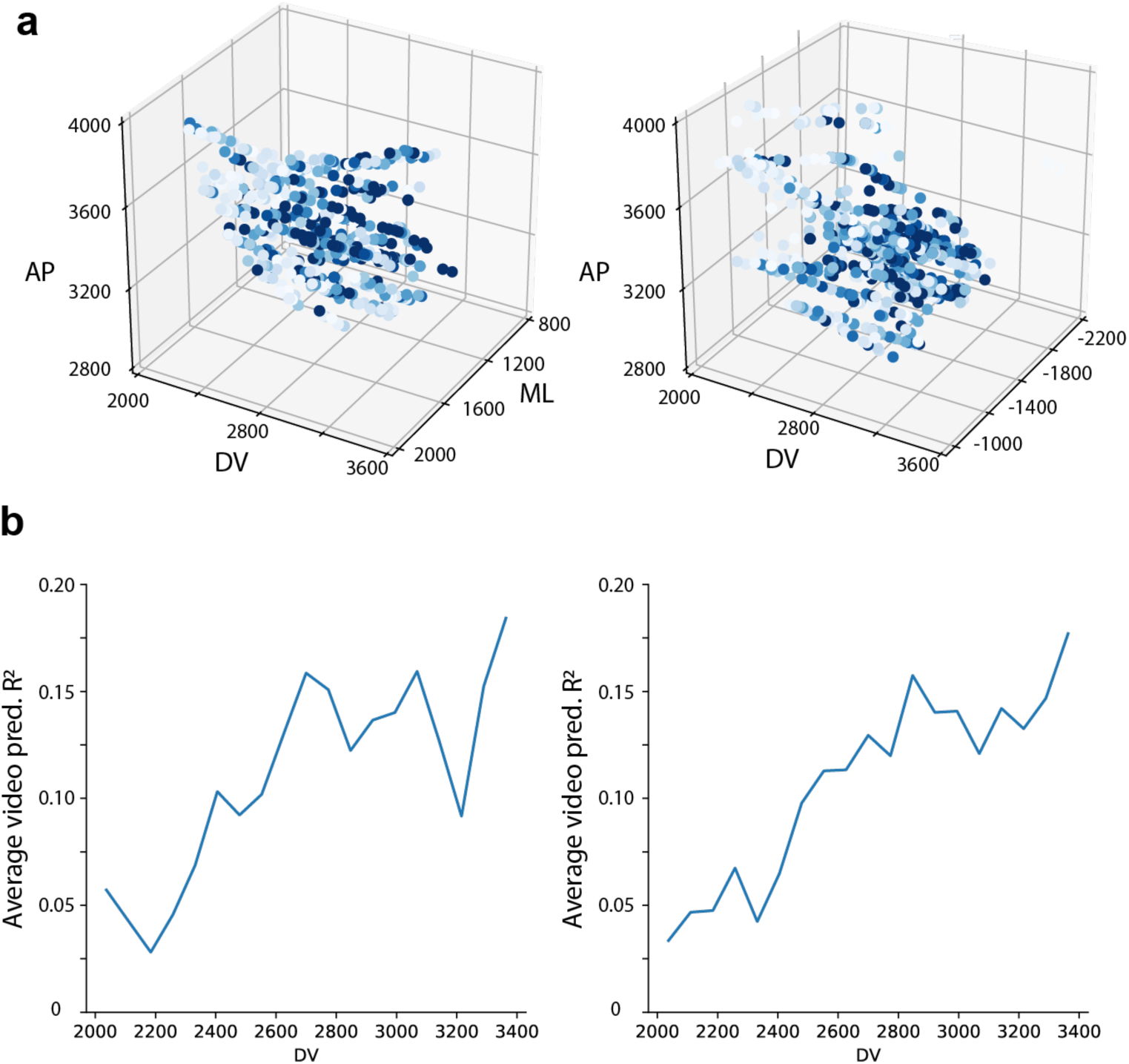
Within region analysis of superior colliculus. **a.** scatter plot of neurons in the superior colliculs in the left hemisphere (left) and right hemisphere (right). Each dot corresponds to a neuron. Color of dot corresponds to variance explained. **b.** average variance explained as a function of dorsal ventral coordinate.

**Figure S6.**
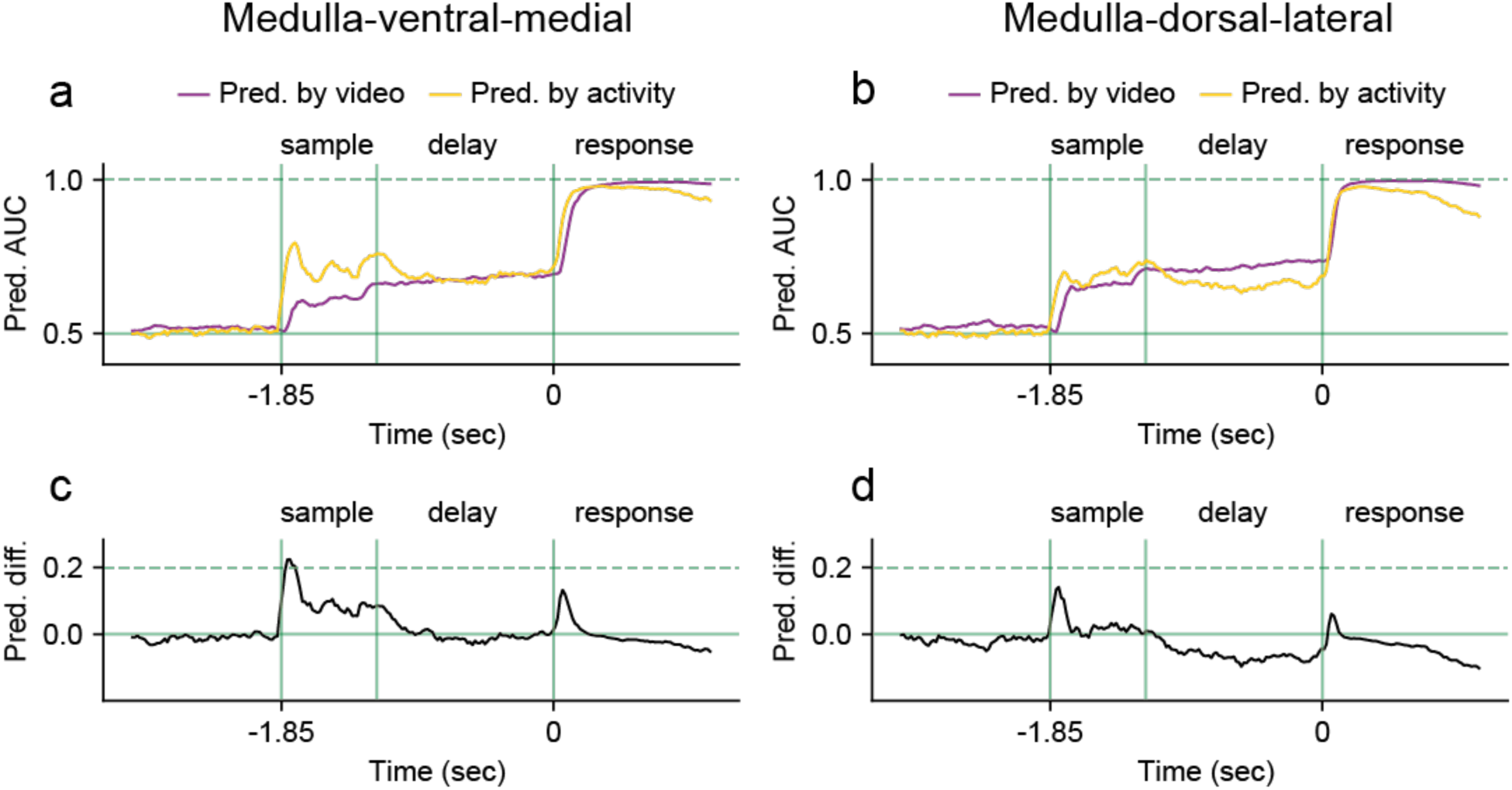
Comparison of time courses of behavior prediction from video or neural activity. **a.** Accuracy of prediction of behavior over time from behavioral videos (purple) and neural activity (orange) for the ventral medial portion of medulla (left) and the dorsal lateral portion of medulla (right). **b.** Difference between the behavioral video prediction and neural activity based prediction as a function of time for the ventral medial portion of medulla (left) and the dorsal lateral portion of medulla (right)

**Figure S7.**
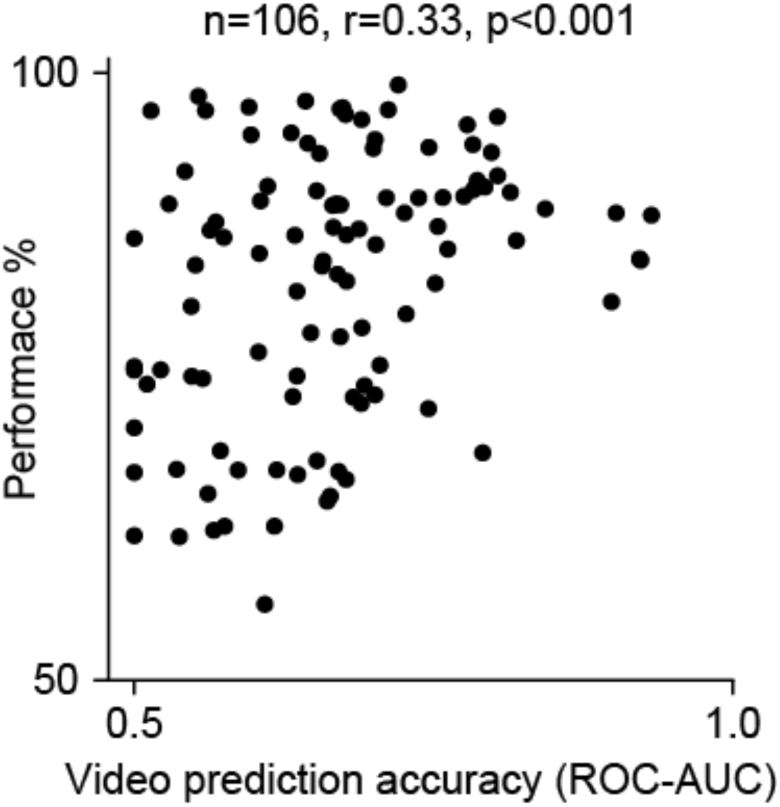
Correlation between predictability of behavior from video and task performance. Each point represents a session. The x axis corresponds to the mean ROC-AUC for predicting behavior directly from video during the delay epoch. Each dot corresponds to a session. The y axis is the behavioral task performance of that session.

